# Longitudinal study of neurochemical, volumetric and behavioral changes in Q140 & BACHD mouse models of Huntington’s disease

**DOI:** 10.1101/2024.02.16.580735

**Authors:** Lori Zacharoff, Ivan Tkac, Alexander Shapiro, Pierre-Gilles Henry, Janet M Dubinsky

## Abstract

Brain metabolites, detectable by magnetic resonance spectroscopy (MRS), have been examined as potential biomarkers in Huntington’s Disease (HD). In this study, the RQ140 and BACHD transgenic mouse models of HD were used to investigate the relative sensitivity of the metabolite profiling and the brain volumetry to characterize mouse HD. Magnetic resonance imaging (MRI) and 1H MRS data were acquired at 9.4 T from the transgenic mice and wild-type littermates every 3 months until death. Brain shrinkage was detectable in striatum of both mouse models at 12 months compared to littermates. In Q140 mice, increases in PCr and Gln occurred in striatum prior to cortex. Myo-inositol was significantly elevated in both regions from an early age. Lac, Ala and PE decreased in Q140 striatum. Tau increased in Q140 cortex. Metabolite changes in the BACHD cortex and striatum were minimal with a striatal decrease in Lac being most prominent, consistent with a dearth of ubiquitin and 1C2 positive aggregates detected in those regions. Binary logistical regression models generated from the Q140 metabolite data were able to predict the presence of disease in the BACHD striatal and previously published R6/2 metabolite data. Thus, neurochemical changes precede volume shrinkage and become potential biomarkers for HD mouse models

Introduction

## Introduction

In the 20 years since discovery of the htt gene, a body of mechanistic knowledge of the myriad physiological consequences of mutant htt expression has emerged from a large variety of experimental models. Some of these mechanisms become apparent in cellular models, some appear only in severe animal models of disease. The difficulty becomes linking these multiple mechanisms to in vivo and most importantly human disease progression. Longitudinal phenotypic behavioral and morphological changes have been studied extensively in the Q140 mouse (Hickey et al., 2008; Lerner et al., 2012). This knock-in mouse has behavioral abnormalities detectable at very young ages, develops neuropathology in parallel, but does not show neurodegenerative changes until the second year of life (Hickey et al., 2008; Lerner et al., 2012). The absence of genetic manipulations beyond the CAG repeat knock-in makes interpretation of any changes that are found uniquely attributable to this mutation.

As a non-invasive technique that can be used in animals from mice to humans, ^1^H MRS provides quantitative information about the chemical composition of brain parenchyma. At high field, concentrations of up to 20 different small molecule metabolites can be obtained in rodents. Previously, we and others have utilized ^1^H MRS in a cross-sectional manner (Jenkins et al., 2000; Tsang et al., 2006; Tkac et al., 2007) to demonstrate that metabolite changes are detectable and useful for monitoring disease and its amelioration by treatments. Longitudinal studies have been performed over the short lifespan of the rapidly progressing R6/2 model or over a few weeks early in disease progression of the extremely mild Q111 model (Tkac et al., 2012; Zacharoff et al., 2012). A longer longitudinal study over the entire lifespan of more slowly progressing mouse models that demonstrates phenotype has been needed to provide detailed information about the progression of these changes. To this end, we examined cortical and striatal regions from the knock in Q140 and BACHD mouse models to yield insights into the onset of various mechanistic consequences of mutant htt expression. The observed metabolite changes are interpreted within the framework of multiple mechanisms contributing to disease progression. If parallel high field studies can be performed in humans, this type of information could be used to inform the timing of mechanistically targeted interventions.

## Methods

### Animals

All experiments were approved by the Institutional Animal Care and Use Committee at the University of Minnesota. The Q140 knock-in mice (Menalled, 2005), homozygous for a 140 CAG repeat knocked into Hdh gene, and their wild-type littermates were bred from pairs obtained from Dr. Michael S. Levine at the David Geffen School of Medicine, UCLA, Los Angeles, USA. These mice were bred on C57BL/6 background strain. Mice hemizygous for a bacterial artificial chromosome containing the full-length human huntingtin gene (BACHD) along with their gene negative littermates were obtained from the same source on an FVB background strain (Gray et al., 2008). Mice were housed in mixed genotype groups, 2-4 per cage, in enriched conditions on a 12:12 light dark cycle and received Purina 5001 pellet food. The Q140 mice (N = 9 ± 1) and littermates (N = 8 ± 1) were scanned in a magnet at 1.5, 3, 6, 9, 12, 15, 18, 21 and 24 months. BACHD mice (N= 8 ± 2) and their littermates (N= 8 ± 2) were scanned at the same intervals beginning at 3 mo. The behavioral tests were performed the week prior to the MRI/MRS scans. Additional backup mice of each genotype were raised as substitutes for mice who died before the end of this longitudinal experiment or when maintenance of 9.4T magnet interrupted the expected schedule. All Q140 and related mice included in this project were scanned at 6 weeks of age and those with high initial levels of brain glutamine, characteristic of a latent portosystemic shunt, (Cudalbu et al., 2013) were excluded from the study.

In both immunohistochemistry and biomarker modeling, tissue and data, respectively, were utilized from a previously published study on the R6/2 model of HD (Zacharoff et al., 2012). R6/2 mice are a severe model of disease, showing behavioral symptoms by 6 weeks and some MRS abnormalities at 4 weeks, the earliest time scanned (Zacharoff et al., 2012).

For the MRI/MRS experiments, spontaneously breathing mice were anesthetized using 1.0 – 1.5% isoflurane in a nitrous oxide and oxygen mixture (1:1). Mice were secured in a cylindrical chamber and their temperature was maintained by temperature controlled circulating water. Respiration was continuously monitored using SAM PC (Small Animal Instruments, Inc., Stony Brook, NY, USA). The total duration an MRI/MRS experiment was 2 hours per animal.

### In vivo magnetic resonance imaging and spectroscopy

All experiments were performed using a horizontal bore 9.4T/31 cm magnet (Varian/Magnex; Oxford UK) equipped with a 15-cm internal diameter gradient coil insert (450 mT/m, 200 µs) with a powerful 2^nd^-order shim set (Resonance Research, Inc., Billerica, MA, USA). The magnet was interfaced to a Varian INOVA console, which was upgraded to the DirectDrive console (Agilent Technologies/Varian, Santa Clara, CA, USA) in a middle of the longitudinal experiment. The MRI/MRS data were acquired using a transmit/receive quadrature surface RF coil with two geometrically decoupled loops (14 mm).

The protocol for collecting MRI/MRS data was very similar to that previously reported (Tkac et al., 2004, 2007; Zacharoff et al., 2012), with a higher level of automatic sequence parameter adjustments for increased throughput of the protocol. Precise position of the volume of interest (VOI) was based on multislice fast spin echo (FSE) images in coronal and sagittal planes (echo train length ETL = 8, echo spacing ESP = 15 ms, TE = 60 ms, matrix = 256 × 256, FOV = 20 mm x 20 mm, slice thickness = 1 mm). The ^1^H MRS data were acquired from VOIs centered in left striatum (VOI = 3.8 – 5.2 µL) and bilateral dorsal cerebral cortex (VOI = 3.8 – 4.8 µL) using ultra-short echo-time STEAM sequence (TE = 2 ms, TR = 5 s) combined with outer volume suppression and VAPOR water suppression (Tkac et al., 1999). The homogeneity of the magnetic field was adjusted automatically using FASTMAP with EPI readout (Gruetter and Tkac, 2000). Brain metabolites were quantified using LCModel with the macromolecule spectrum included in the basis set (Provencher, 1993; Pfeuffer et al., 1999; Tkac et al., 2004). The unsuppressed water signal was used as an internal concentration reference assuming 80% brain water content. The concentrations of the following brain metabolites were routinely quantified in both brain regions with Cramér-Rao lower bound (CRLB) below 15%: creatine (Cr), phosphocreatine (PCr), γ-aminobutyric acid (GABA), glutamine (Gln), glutamate (Glu), *myo*-inositol (mIns, Ins), lactate (Lac), N-acetylaspartate (NAA), taurine (Tau) and the sum of glycerophosphocholine + phosphocholine (GPC+PC). Glucose (Glc), glutathione (GSH) and phosphoethanolamine (PE) were quantified with CRLB < 25%, ascorbate (Asc), N-acetylaspartylglutamate (NAAG), alanine (Ala) in striatum and aspartate (Asp) in cortex were quantified with CRLB < 35%. Total creatine (Cr+PCr) and the macromolecule (MM) content were also reported.

For the volumetric measurements, multislice coronal MR images, acquired using a high resolution FSE imaging sequence (ETL = 16, ESP = 8 ms, TE = 64 ms, slice thickness = 0.3 mm, in-plane resolution = 100 µm x 100 µm, number of slices = 24, number of averages = 16, total measuring time 22 min), were obtained from the rostral pole of the frontal cortex to the caudal pole of the occipital cortex. Images were evaluated by user guided segmentation (Amira, Visage Imaging, San Diego, CA, USA) to determine regional brain volume changes in whole brain, cortex, striatum, lateral ventricles and third ventricle. The boundaries of the cortex were defined ventrally by the corpus callosum and lateral ventricles. The striatum was defined by the corpus callosum dorsally and laterally, by the lateral ventricles medially and by the anterior commissure ventrally. Anterior brain measurements indicating overall brain size were summed from all structures in all image planes, including ventricles.

### Climbing Assay

During daylight hours, individual mice were placed in an upside-down wire mesh pencil holder for 5 min and videotaped (Hickey et al., 2005). At least 2 independent, blinded observers scored the videos for climbing and rearing behaviors, calculating the total time spent, the latency to onset and the number of instances of climbing and rearing. Climbing was defined as all 4 paws off of the ground and on the wire mesh. Rearing was defined as 2 - 3 paws off of the ground, not necessarily on the wire mesh.

### Immunohistochemistry

Brains were harvested from rapidly sacrificed mice, embedded in OCT, frozen in liquid isopentane at −70°C and subsequently stored at that temperature. For immunocytochemistry series of 30µm slices were fixed in 4% paraformaldehyde, rinsed, pretreated sequentially with formic acid and hydrogen peroxide, and exposed to primary antibody for 24hr. For 1C2-immunoreactivity, sections were treated with formic acid for 3 minutes at room temperature prior to fixation to expose the contents of the intracellular inclusions to this antibody (Herndon et al., 2009). After rinsing, slices were exposed to secondary antibodies conjugated to HRP (FD Neuro Technologies) and subsequently reacted with DAB. The primary antibodies used were a polyclonal against ubiquitin (Dako 1:400) (Davies et al., 1997), and the monoclonal 1C2 (Millipore 1:5,000)(Trottier et al., 1995).

### Statistics

Two-way ANOVAs followed by Bonferroni post-tests (GraphPad Prism, v. 6.0) were used for a statistical evaluation of longitudinal metabolic, volumetric and behavioral data measured from striatum and cerebral cortex of all mice. Binary logistical regression modeling was performed in Minitab, version 16. ROC analysis was performed by GraphPad Prism, v. 6.0.

## Results

### Longitudinal changes in neurochemical profiles of Q140 mice

*In vivo* ^1^H MR spectra from the striatum and cerebral cortex of Q140 mice and littermate controls were acquired at 6 and 12 weeks of age and subsequently every 3 months over their 2-year-long lifespan. Maintaining consistent, high quality spectra (Fig. 1) over the two year duration of this project (> 300 spectra) was critical for reliable metabolite quantification. Longitudinal trajectories of a broad range of cortical and striatal metabolites reveal variable patterns of change among Q140 metabolites compared to littermates (Fig. 2, Supplementary Fig. 1). Complete cortical and striatal neurochemical profiles quantified from all mice and littermate controls at all ages can be found in the Supplementary Table 1. In general, metabolic changes between Q140 and littermate mice were more prominent in striatum relative to cortex. The most significant differences in striatal metabolite levels between Q140 and littermate mice were observed for Gln, PCr, and Cr+PCr. Striatal concentrations of these metabolites in Q140 mice progressively increased with age and a strong correlation (R = 0.80, p < 0.0001) was found between total creatine and Gln (Fig. 3). Concentrations of Ala, Lac and PE progressively decreased in Q140 mice relative to littermate littermates (Fig. 2). In addition, the two-way ANOVA tests revealed small, but significant differences between genotypes for Glu, Ins, Tau and macromolecules (MM). On the other hand, concentrations of NAA and total cholines (GPC+PC) were extremely stable over the 2-year period without any difference between genotypes. Concentrations of GABA, Glc and GSH in striatum did not show age or genotype related differences. Asc and NAAG were not consistently quantified in all age groups and striatal Asp could not be detected.

**Fig. 1.**
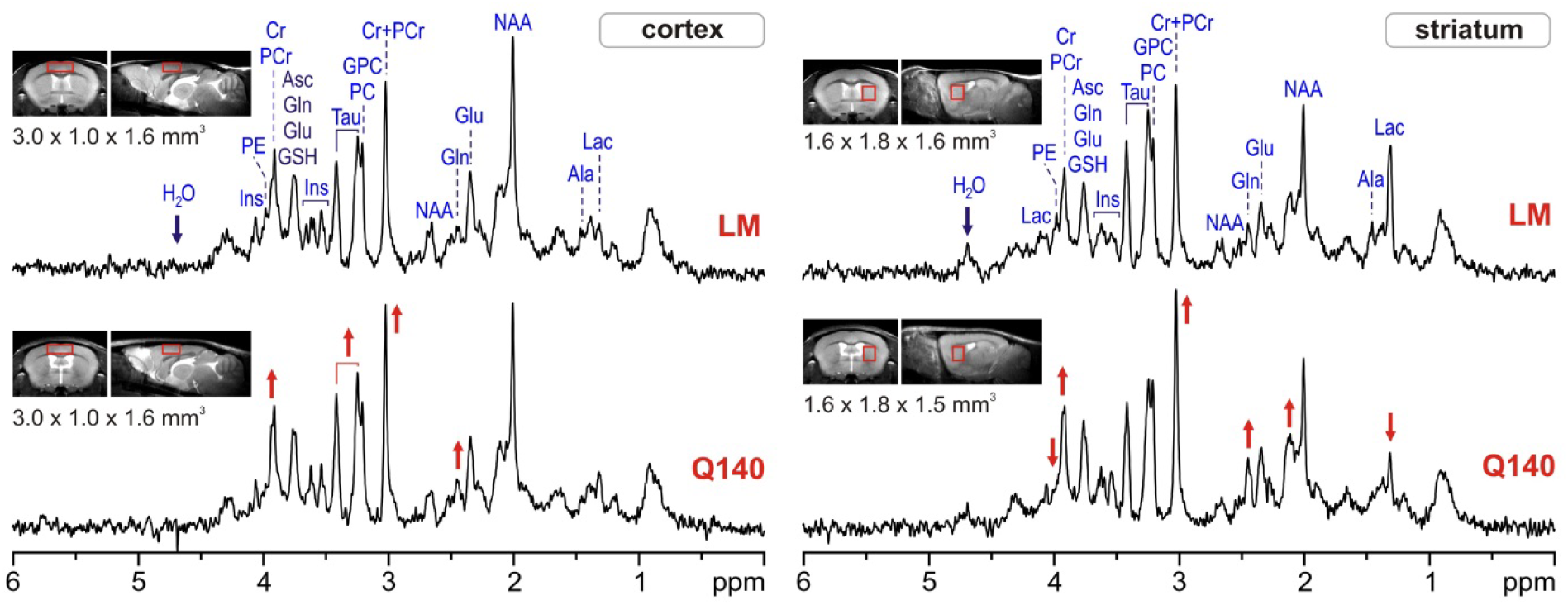
1H MRS spectra for cortical and striatal voxels (indicated in inserts) from Q140 and LM mice. Peaks of prominent metabolites are identified. Red arrows indicate changes observed over the lifespan of the Q140 mice.

**Fig. 2.**
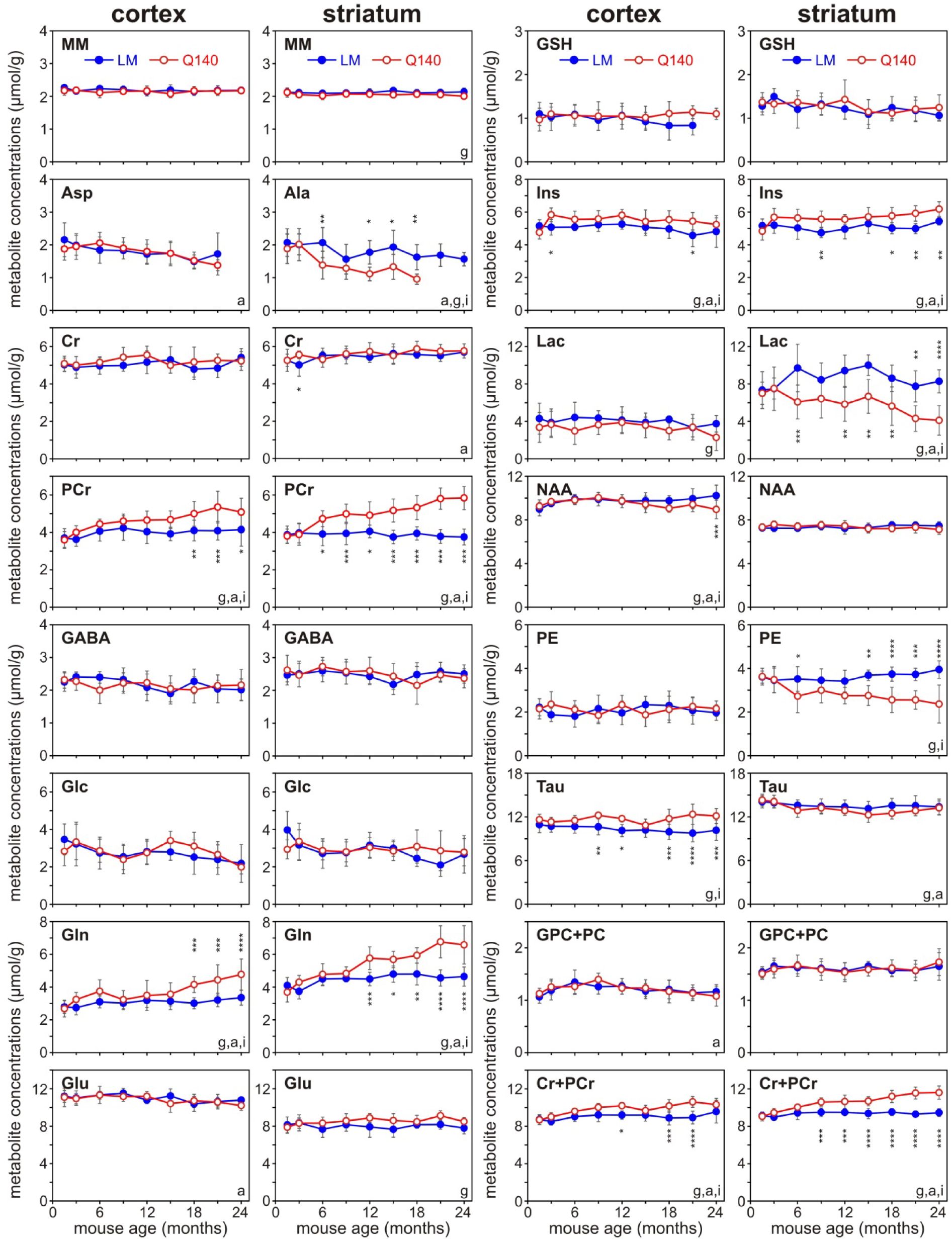
Longitudinal changes in 16 striatal and cortical metabolites over the lifespan of Q140 (open red circles) and littermate (closed blue circles) mice. Data are mean ± standard deviation from 8 or more mice of each genotype per time point. Significance of two way ANOVAs main effects for each metabolite comparing the ability of genotypes, age and their interactions to explain variability at the p<0.01 level or greater shown by g, a, or i, respectively. *, **, *** indicate p<0.05, p<0.01, p<0.001, respectively for individual time points in Bonferonni corrected post-tests.

**Fig. 3.**
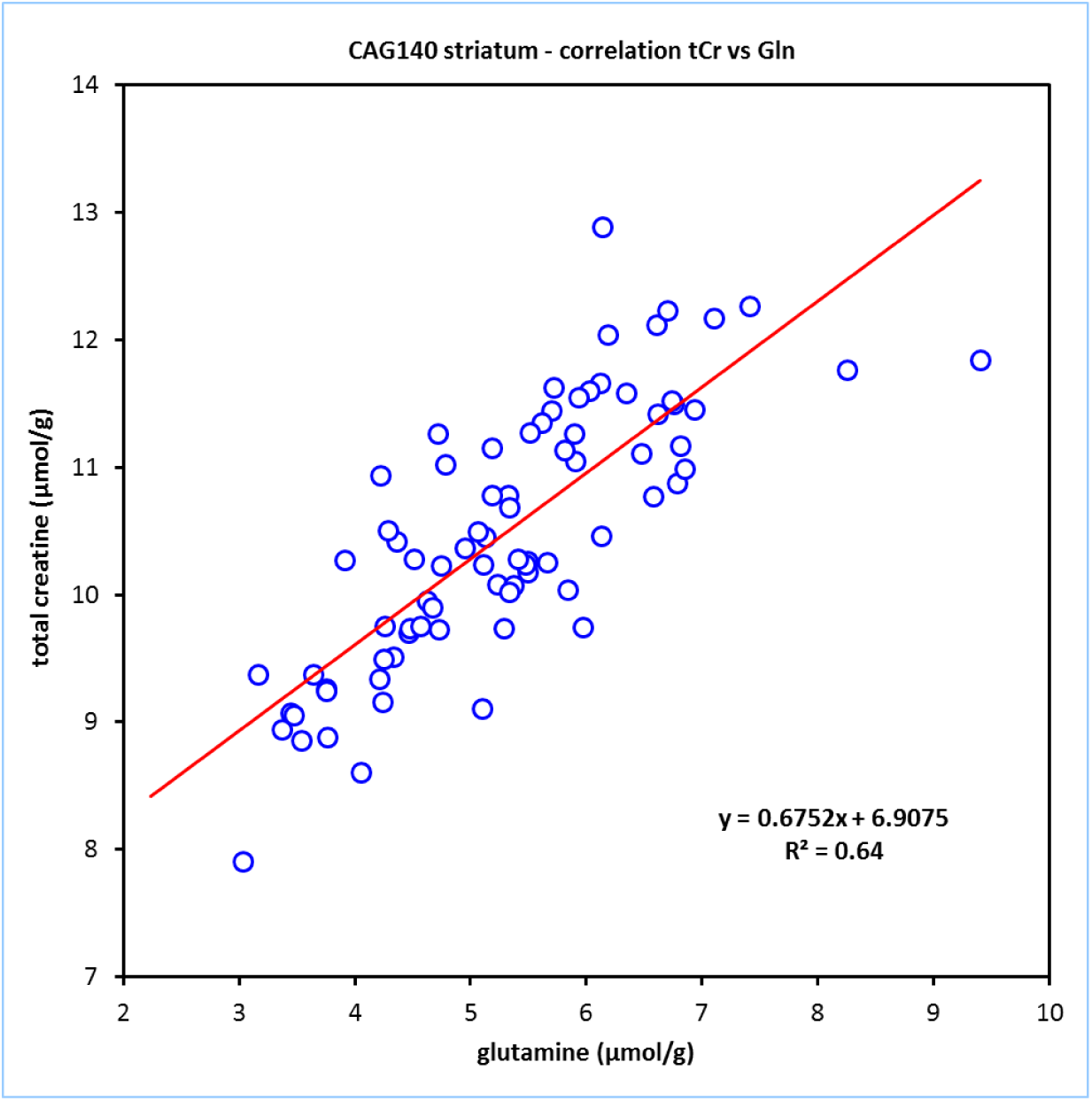
Correlation between the detected concentrations of glutamate and total creatine (PCr+Cr) over all ages.

The most significant concentration changes in the cerebral cortex between Q140 and littermate mice were observed for Gln, PCr and Cr+PCr, similarly to striatum. However, the amplitude of these changes were on average smaller than in the striatum and were only observed in older age groups (Fig. 2). In contrast to the striatal data, cortical Tau concentrations were significantly lower in Q140 mice relative to littermate controls. Small, but significant decreases in NAA were observed in Q140 mice at the oldest ages. Differences between genotypes were much smaller for cortical Lac and not observable for PE. The concentrations of Asc (not shown) and Asp decreased and NAAG increased (not shown) with age, but genotype differences were not observed.

### Longitudinal changes in neurochemical profiles of BACHD mice

Metabolite changes in the BACHD (Supplementary Fig. 2) have also not been as prominent as in the R6/2 or Q140 mice. In striatum, Lac, GABA and Ala decreased in BACHDs compared to littermates, while Glc, PCr and mIns may be increased. In BACHD cortex, Lac also decreased, and Glc and NAAG increased compared to littermates. Very few of these changes reached significance in post-tests, but the 2 way ANOVAs indicated that genotype contributed significantly to the variation for these metabolites. Age dependent changes were observed in many other striatal and cortical metabolites. For the most part, the observed differences were not as great between BACHDs and their controls as were comparable changes in the Q140s with respect to their littermates. Only the decrease in striatal Lac concentration appeared to be progressive as the BACHD mice aged.

### Brain volumetric measurements and body weight

Robust tissue segmentation was readily obtainable from the high resolution, high contrast FSE images (100 x 100 x 300 µm^3^, Fig. 4 top and middle). Highly significant differences in striatal volumes were observed between Q140 and littermate mice (Fig. 4 bottom). The decreased striatal volume in Q140 mice relative to WT controls was first detected at 3 months of age. Q140 striatal volume remained stable until 15 mo when it began to decline further. Differences in cortical and anterior brain volumes between Q140 and littermate mice were observed only at older ages. No differences in lateral or 3^rd^ ventricle volumes or body weight were observed between genotypes (Fig. 4 bottom, Supplementary Fig. 3).

**Fig. 4.**
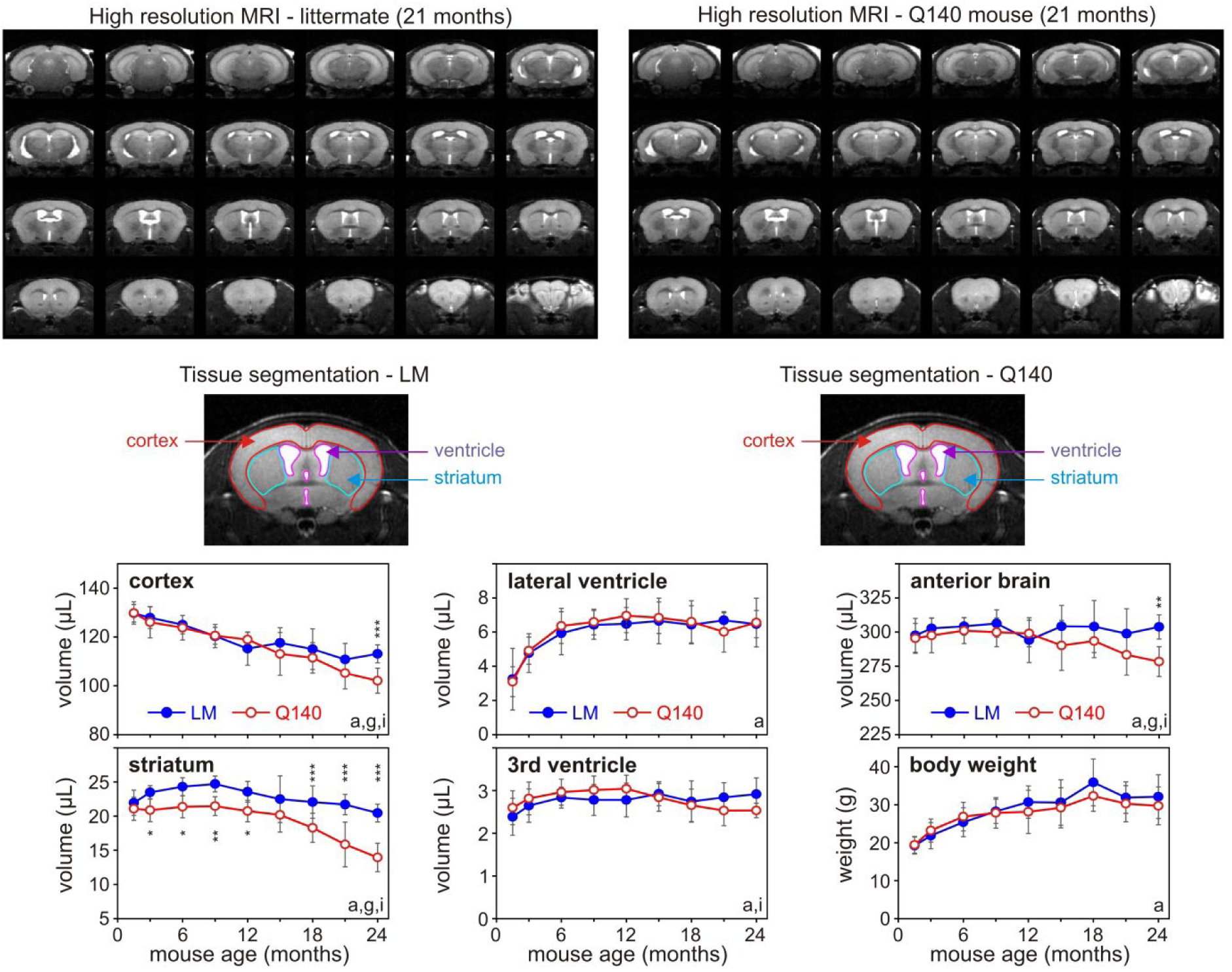
Analysis of regional volume changes measured in coronal images from Q140 and littermate mice. Top: RARE images from typical 21 mo old Q140 and littermate mice. Middle: illustration of the manual segmentation applied to volumetric measures for striatum, cortex, lateral and third ventricles and entire anterior brain. Bottom graphs: Longitudinal changes in regional volumes. Statistical results coded as in Fig. 2.

As in the Q140s, the BACHD striatal volume was consistently lower than that of littermates from the age of 3 mo (Supplementary Fig. 4). Regional brain volumes of BACHD mice (cortex and anterior brain) were comparable to or slightly below that of littermates throughout the lifespan, but more noticeable in the last months of life. No differences were observed in the ventricular volumes by genotype.

All mice gained weight normally as they aged (Supplementary Fig. 3). However, BACHD mice gained more weight than their littermates, as previously noted for this strain (Gray et al., 2008).

### Behavioral studies

The videotaped climbing tests of mice were analyzed and six different behavioral criteria were evaluated (Fig. 5). The assessment of instances of rearing was the only behavioral test where progressive differences between Q140 mice and littermate controls were observed. Climbing behavior did not differentiate Q140 mice littermate controls. All mice accommodated to the climbing chamber and explored it less with the regular repeated procedure, as evidenced by the decreased numbers of climbs or rears. When a 12 month old Q140 littermate mouse who had not previously been tested was put into the apparatus, the latency to rear or climb was less and the instances of rearing and climbing were greater than the mice in the longitudinal study.

**Fig. 5.**
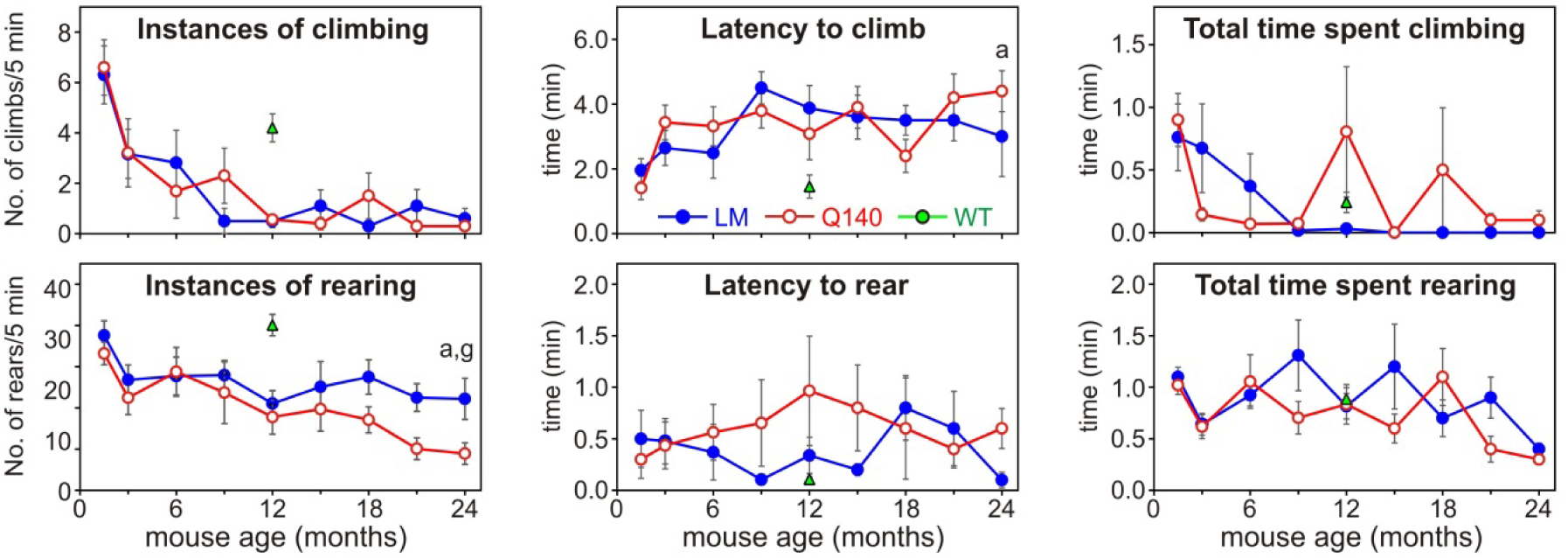
Longitudinal changes in climbing assay behavior over the Q140 (red open circles) and littermate (blue closed circles) lifespan. At 12 months, naïve littermate mice, not previously exposed to the climbing chamber were tested at a single time point (green triangle). Statistical results coded as in Fig. 2.

The BACHD mice also appeared to accommodate to the climbing assay, more so than their littermate controls (Supplementary Fig. 5). While these behavioral measures did distinguish disease, they could easily be explained by the weight gain in the BACHD mice compared to littermates.

No clasping behavior was seen at any age in Q140, BACHD or their littermates.

### Immunohistochemistry

Since so few metabolic changes were detected in the BACHD mice throughout their lifespan, we sought other measures to determine if disease was progressing. BACHD mouse brains were examined for the presence of huntingtin aggregates in both striatum and cortex upon sacrifice at 24 mo. R6/2 brains at 12 weeks of age were examined as positive controls. Tissue from Q140 brains were not available. Few mutant huntingtin aggregates were encountered in striatum and cortex of BACHD mice compared to R6/2 mice. Immunostaining for Mab 1C2 which binds to the toxic polyglutamine conformation (Trottier et al., 1995) (Fig. 6, A-H) and a polyclonal antibody that recognizes ubiquitin (I-P) revealed multiple punctate structures in striatum and cortex of 16 wk R6/2 brains (Fig. 6 C, G, K, O) but few puncta in 24 month BACHD brains (Fig. 6 A, E, I, M). The majority of fields in the BACHD brains had no discernible immunoreactivity. This confirmed that disease progression was extremely slow in the BACHD cohort.

**Fig. 6.**
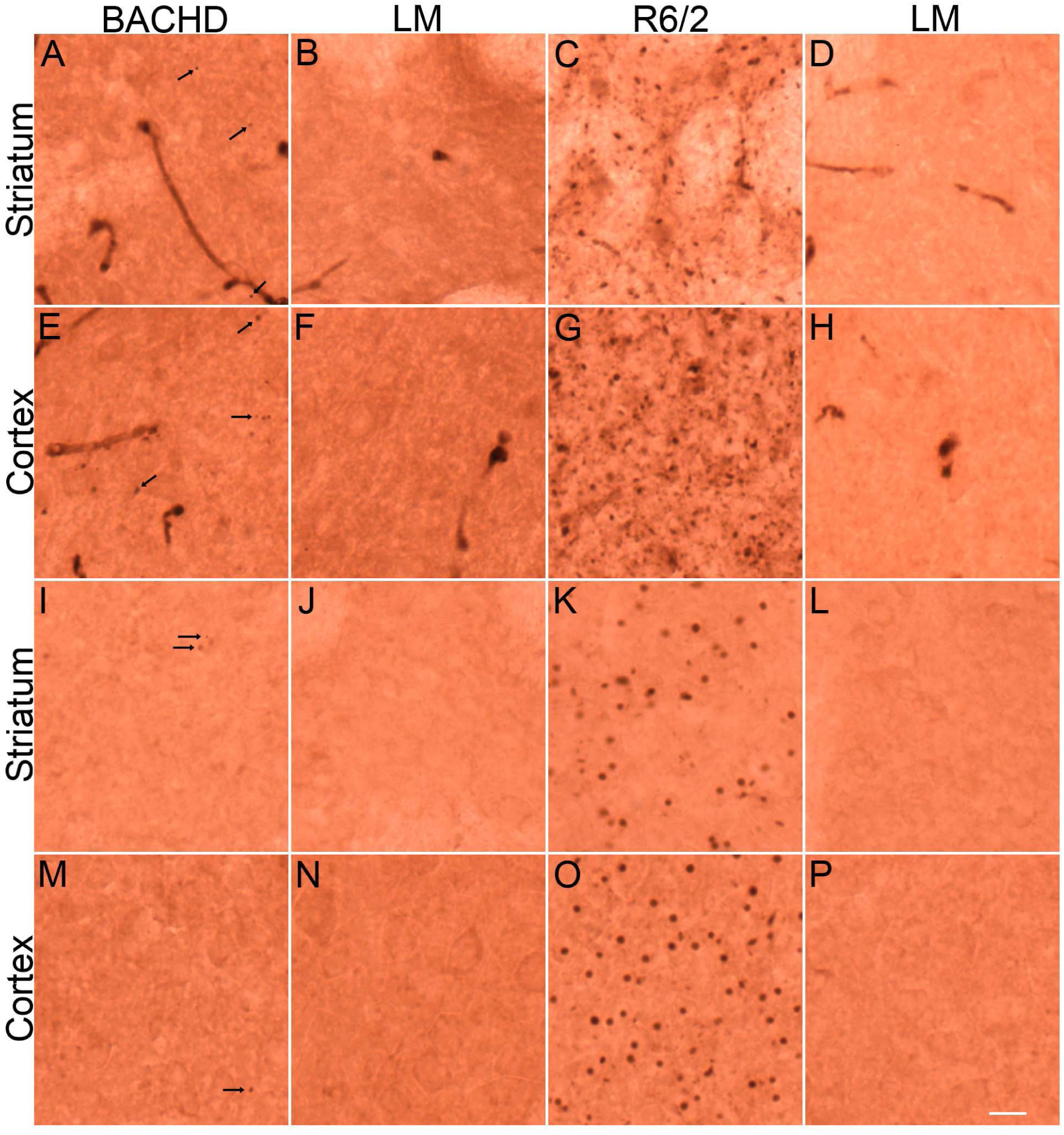
Immunohistochemistry of intracellular aggregates in BACHD brains revealed very few puncta compared to R6/2 brains. Immunostaining for Mab 1C2 (A-H) and immunostaining for ubiquitin (I-P) in striatum (A-D, I-L) and cortex (E-H, M-P) of BACHD (A, E, I, M) and R6/2 (C, G, K, O) brains. Staining from littermate control brains corresponding to each HD mouse type are shown to the right of that HD mouse (B, F, J, N, D, H, L, P). All puncta in A, I and M are labeled with arrows. Additional puncta in E are not labeled. Scale bar = 10μm.

### Biomarker Construction

We sought a method that would combine metabolites and anatomical changes into a model to predict disease across the R6/2, Q140 and BACHD mouse strains. Binary logistic regression was chosen as a method commonly used in the clinical literature for its ability to deliver a binary outcome of diseased or not from multiple input variables (Grant et al., 2018). Binary logistic regression models were developed using the longitudinal data from the striatum and cortex of Q140 mice and their wild type littermates. An initial model included 10 metabolites, weight, sex and age (Table 1, model 1). This yielded a good log-likelihood value indicative of model fit (the higher the value, the better the fit). While none of the predictor variables were individually highly significant, together they effectively distinguished among Q140 mice and their littermates. The model returned a probability value that each mouse was normal (*p*=1, normal; *p*=0, diseased; Fig. 7a,b). These calls clearly distinguished Q140 mice from littermates at 6 mo of age. Accuracy of the calls increased with age. As some of the confidence intervals of these variables were broad, we examined models with fewer variables. Models with single predictor metabolites had the narrowest confidence intervals and lowest *p* values (Table 1 model 2), as would be expected from the changes seen in individual metabolites (Fig. 2, Supplementary Fig. 2), but these did not incorporate information from other measures. In a model combining age and Cr, PCr, Gln, and Lac, each predictor’s contribution was highly significant and had tight confidence intervals (Table 1, model 3). Further addition of anatomical volume measurements improved the overall fit, while decreasing the significance of each predictor (Table 1, model 4). We did not attempt to optimize models further.

**Fig. 7.**
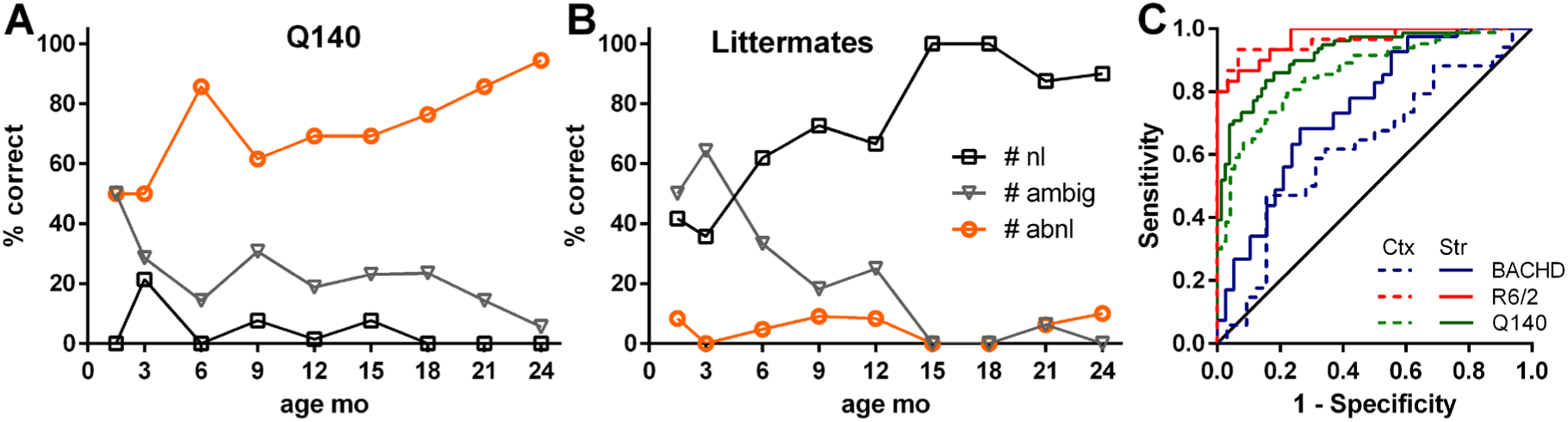
Binary logistic regression models distinguish diseased from non-diseased mice. A, B) Probability calls for Q140 (A) and littermate mice (B) using Model 1 from Table 1. Thresholds applied were: p ≥ 0.75 plotted as normal (nl, black squares), 0.25 < p < 0.75 plotted as ambiguous (ambig, grey triangles), p ≤ 0.25 plotted as abnormal (abnl, orange circles). C) Receiver-operator curve analysis of probability calls from Model 3, Table 1 made from Q140 data applied to 12-24 mo BACHD data and previously published R6/2 data (Zacharoff et al., 2012). Curves represent R6/2 (red), BACHD (blue) and Q140 (green) data for striatum (solid line) and cortex (dashed line). See Table 2 for AUC and associated statistics.

**Table 1:**
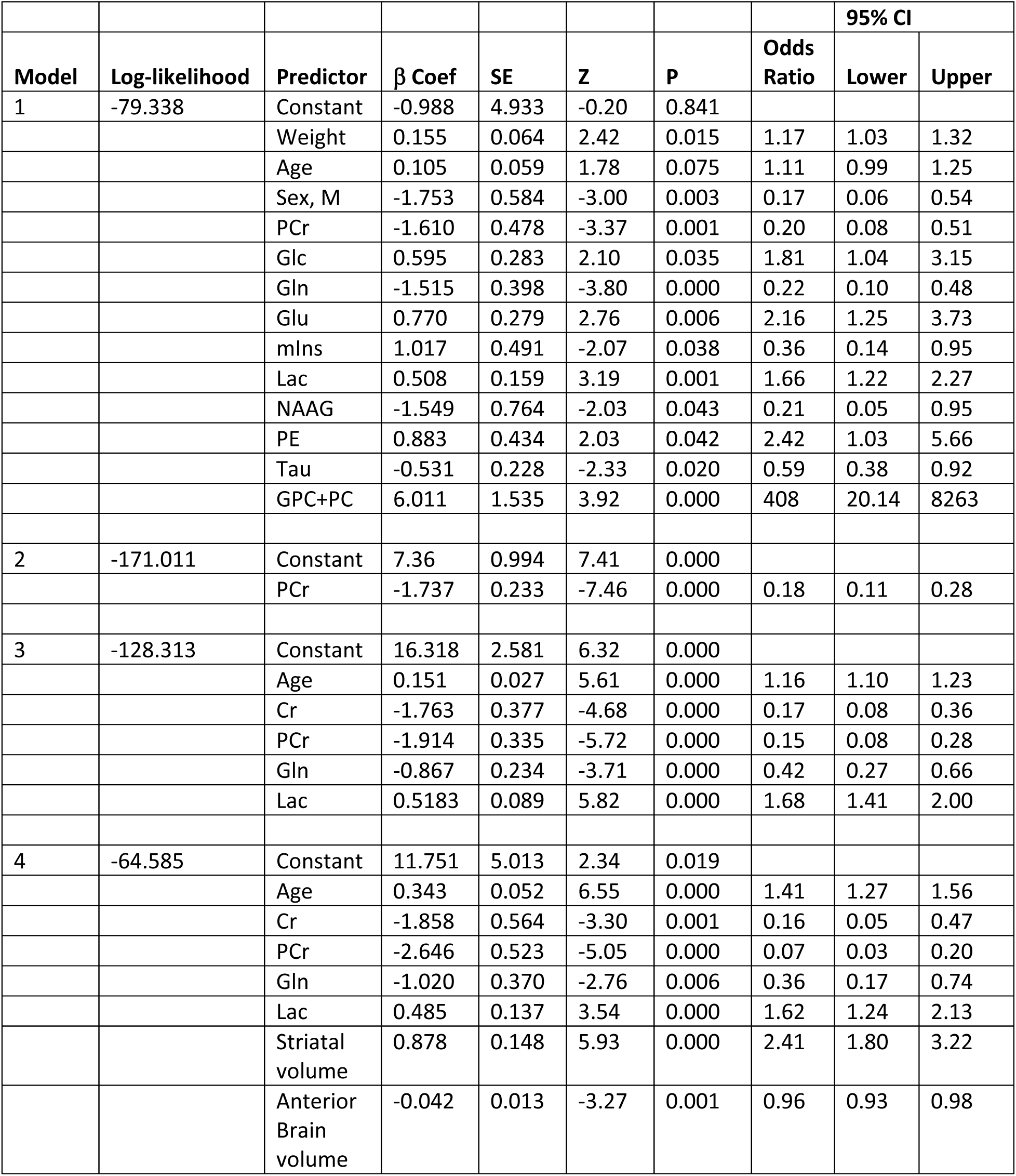
Table of Logistic Regression Models.

Instead, we tested whether a reduced, combinatorial model made from Q140 data (Fig. 2; Table 1, model 3; Fig. 7) could predict the presence of the disease among the BACHD mice, ages 12-24 mo (Supplementary Figs. 2, 4, 5), and R6/2 mice (Zacharoff et al., 2012). A receiver-operator curve (ROC) analysis of the probabilities of normality returned for all mice demonstrated that even a non-optimized binary logistic regression model clearly distinguishes the HD mice by known severity of disease (Fig. 7c). Areas under the ROC curve are significant for all mice and regions except the BACHD cortex (Table 2).

**Table 2.**
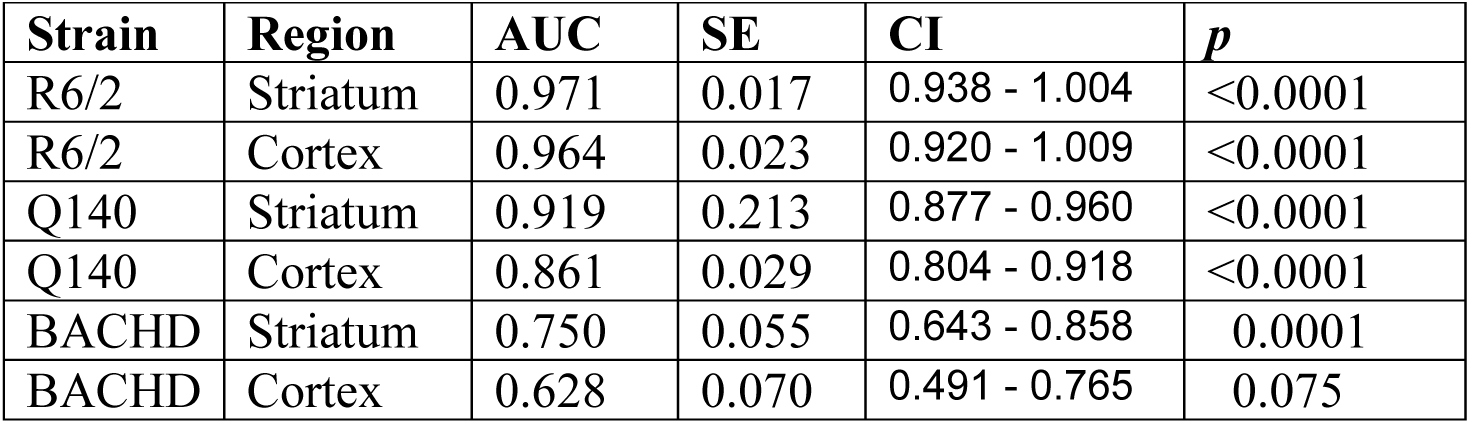
ROC analysis (see Fig. 7)

| **Strain** | **Region** | **AUC** | **SE** | **CI** | ***p*** |
| --- | --- | --- | --- | --- | --- |
| R6/2 | Striatum | 0.971 | 0.017 | 0.938 - 1.004 | <0.0001 |
| R6/2 | Cortex | 0.964 | 0.023 | 0.920 - 1.009 | <0.0001 |
| Q140 | Striatum | 0.919 | 0.213 | 0.877 - 0.960 | <0.0001 |
| Q140 | Cortex | 0.861 | 0.029 | 0.804 - 0.918 | <0.0001 |
| BACHD | Striatum | 0.750 | 0.055 | 0.643 - 0.858 | 0.0001 |
| BACHD | Cortex | 0.628 | 0.070 | 0.491 - 0.765 | 0.075 |

## Discussion

This longitudinal study provides comprehensive results on neurochemical and volumetric changes in the Q140 and BACHD mouse models of HD at regularly spaced intervals over their entire 2 year lifespan. Volumetric and neurochemical changes were expected to be primarily localized in the striatum, mimicking the phenotypes observed in human HD (Tang and Feigin, 2012; Clabough, 2013). Indeed, changes in both measurements in Q140 and BACHD mice were more pronounced in the striatum than in the cortex (Figs. 2, 4 and Supplementary Figs. 2, 4). Common to both strains were a decrease in Lac and Ala and an increase in PCr and mIns in the striatum. Striatal volume also failed to increase from 3 to 12 months and declined thereafter. Behaviorally, both strains demonstrated accommodation to the climbing assay and failed to show clasping behaviors, consistent with these strains mimicking early disease stages. In BACHDs, the progressive weight gain may also have contributed to a failure to climb. We interpret these changes in the context of the onset of pathologies and behaviors characteristic of these slowly progressive mouse models of HD.

### Volumetric data

Early in life, striatal volume in Q140 and BACHD mice remained stable, while that in littermates increased, suggesting a possible reduced rate of brain growth and development rather than brain atrophy (Nopoulos et al., 2011; van der Plas et al., 2020). The failure of the striatum to grow may be attributable to the inability to form new dendritic spines in a BDNF-depleted environment. A decreased spine or synapse density and less complex arbors characterize both rapidly progressing (R6/2, Q175) and more slowly progressing (R6/1, Q89, YAC72, YAC128, Q140) models once disease is manifest (Hodgson et al., 1999; Guidetti et al., 2001; Slow et al., 2003; Spires et al., 2004; Lerner et al., 2012; Zarate et al., 2023). At younger ages, cortical spine density was unchanged and dendritic abnormalities were subtle in symptomatic R6/1 mice at 20wk, but pronounced in R6/2 striatum at 4 wk, an age largely considered presymptomatic (Nithianantharajah et al., 2009; Heck et al., 2012). Selective ablation of BDNF in an otherwise normal cortico-striatal pathway mimics the morphological changes observed in many HD mouse models, namely somal shrinkage, thinning and shortening of dendrites and loss of spine density, observable as early as postnatal day 35 (Baquet et al., 2004). In hippocampal slices from 8 wk old Q140 mice, theta burst stimulation fails to induce LTP and increase phalloidin staining of actin in new spines, effects that were ameliorated by treatment with an ampakine that mimics BDNF (Simmons et al., 2009). Thus it is likely that spine abnormalities may be present in young Q140 and BACHD mice, but this needs to be systematically examined with stereological techniques.

Similarly, axonal function and morphology may be altered in HD brains. In Q140 mice, striatal volume was significantly decreased by 12 mo but the somal size distribution was unchanged at ages over 20 mo (Fig. 4 and (Lerner et al., 2012)). In BACHD mice at 15 mo, the decreased striatal volume becomes more pronounced. At this advanced age, shrinkage of the cortex and corpus callosum were also observed, suggesting the loss of axonal projections accompanies later stages of disease (Lerner et al., 2012). Small htt aggregates have been found in neuropil, axons and nerve terminals in 2-4 mo Q140 mice, suggesting that early behavioral deficits may be influenced more by axonal and synaptic dysfunction than by degeneration (Hickey et al., 2008). Both dark cell neuronal degeneration and axonal degeneration in striatum have been observed at the EM level in 14 mo old CAG150 mice, a similar, slowly progressing knock in model of HD (Yu et al., 2003). Loss of axons and white matter is observed in presymptomatic HD gene carriers in both cortex and the internal capsules (Weaver et al., 2009; Rosas et al., 2010). Since in mice the internal capsule traverses the striatal voxel scanned in our study, loss of a proportion of these axons cannot be ruled out.

### 1H MRS data

Despite this loss of volume, striatal NAA levels remained constant over the entire lifespan in both mouse models. In contrast, cortical volume and NAA concentration remained stable over most of the lifetime, decreasing in parallel beginning at 15 mo but becoming significant only at 24 mo in the Q140s. The absence of changes in striatal NAA was puzzling. NAA is considered a marker of neuronal health since it is synthesized in neurons, degraded in oligodendrocytes, not found in astrocytes, and often decreases in degenerative diseases (Satrustegui et al., 2007; Duarte et al., 2012). The unchanged striatal levels of NAA are consistent with the relatively stable levels of Glu, as both these metabolites primarily share the neuronal compartment (Duarte et al., 2012). A more nuanced way to interpret NAA may be as a marker of neuronal function since NAA levels can recover after transient ischemia (Duarte et al., 2012). Behavioral assays used here were not sensitive enough to detect motor decline. Considering NAA is synthesized in neuronal mitochondria and degraded in oligodendrocytes, NAA decreases may signal mitochondrial dysfunction, demyelination or both (Satrustegui et al., 2007). Interpretation of Q140 NAA awaits replication with more exhaustive behavioral and anatomical measures.

PCr increased progressively beginning around 3 - 6 mo (Fig. 2, Supplementary Fig. 2) in striatum of both strains. The increase in PCr appears to follow the onset of the reduction in CK expression between 2 and 4 mo in Q140 mice (Kim et al., 2010). Decreased CK activity may create a bottle neck, reducing the availability of this high energy compound (Zhang et al., 2010). Similarly creatine kinase protein expression was reduced at 3 wk in R6/2 and PCr levels began to climb in that model at 4 wk (Kim et al., 2010; Zacharoff et al., 2012). Creatine regulation is governed by the synthetic enzymes arginine-glycine amidino-transferase (AGAT) and guanidinoacetate methyltransferase (GAMT) that are expressed heterogeneously across different neural cell types (Andres et al., 2008; Braissant and Henry, 2008; Braissant et al., 2010; Béard and Braissant, 2010). The SLC6A8 transporter moves Cr across the blood-brain-barrier and creatine and its precursors between neural cells (Tachikawa et al., 2008; Béard and Braissant, 2010). Changes in mRNA levels and/or protein expression of GAMT and CK, but not AGAT or SLC6A8, were observed in striatum and frontal cortex of R6/2 mice (Mochel et al., 2012). These have not been explored for Q140 or BACHD mice.

Lac and the biochemically linked Ala decreases in the striatum of both strains may reflect glucose hypometabolism, a finding consistent across multiple mouse models and premanifest and manifest human disease (Berent et al., 1988; Antonini et al., 1996; von Horsten et al., 2003; Wang et al., 2005; Ciarmiello et al., 2006; Cepeda-Prado et al., 2012; Li et al., 2012). Decreased glucose consumption and subsequent lactate production would be consistent with the interpretation that the progressively increasing PCr/Cr observed throughout the Q140 lifespan reflects a decreased energy demand. Given that Lac levels correlate with increased brain activity (Mangia et al., 2007; Lin et al., 2012; Schaller et al., 2013), these data suggest a decreased energy demand from the mutant htt sensitive inhibitory medium spiny striatal neurons which create the majority of striatal metabolic demand (McCasland and Hibbard, 1997; Dubinsky, 2009, 2017; Cepeda-Prado et al., 2012). The decreased Lac observed in the current study contrasts with the marked increase in Lac consistently observed in the R6/2 (Tkac et al., 2007; Zacharoff et al., 2012). Similarly, glial glycolytic flux has been suggested to increase in human HD (Powers et al., 2007). These differences may indicate that metabolic compromise and/or dysfunction as reflected by Lac levels differs between slowly and rapidly progressing mouse models or human disease stages. Considering that the neuropathology of mutant htt varies depending upon the protein context (Yu et al., 2003), the metabolic responses to the presence of mutant htt may also vary depending upon context. Decreased striatal levels of Ala paralleled those of Lac, as these metabolites are in dynamic exchange though pyruvate (Wolfe et al., 1988).

Gln resides primarily in the astroglial compartment and is considered to be a glial marker (Norenberg and Martinez-Hernandez, 1979; Daikhin and Yudkoff, 2000; Hertz, 2004; McKenna, 2007). Astrogliosis was not visible by immunostaining at 4 mo in the Q140 cortex or striatum but became extensive by 12 mo in cortex and 24 mo in striatum (Hickey et al., 2008). A different astrogliosis progression in this cohort of Q140 mice cannot be ruled out as no animals were sacrificed for neuroanatomical investigation. The increase in striatal Gln may also reflect an additional role of Gln as an osmoregulatory (Thurston et al., 1983; Heilig et al., 1989; Verbalis, 2010). Osmoregulators substitute for Na in hypernatremia, a process ultimately dependent on energy metabolism and the Na/K ATPase (Thurston et al., 1983; Heilig et al., 1989). If osmotic load contributes to morbidity in HD, it is likely to be observed first in the striatum, a region susceptible to energy impairments (Dubinsky, 2009). An osmoregulatory role for Gln is also supported by the observation of its rise in the R6/2 model (Tkac et al., 2007; Zacharoff et al., 2012) where astrogliosis is not observed (Yu et al., 2003)(Dubinsky, unpublished observations).

In Q140 mice, the step increase in mIns at 3 mo in both regions occurred too early to support a role of mIns as a precursor of increased synthesis of inositol containing phospholipids in gliosis, as has been hypothesized (Duarte et al., 2012). This step increase is small (11%) and does not increase further at ages when gliosis has been documented. The small increase may reflect establishment of a new regulated steady state rather than a progressively evolving pathogenic process. mIns was also elevated in the BACHD striatum prior to the appearance of aggregates. Increased mIns prevents autophagy stimulation in HDQ74-expressing cells (Williams et al., 2008). Increased inositol and subsequently IP3 activate an IP3-receptor dependent Ca^2+^ release from the ER blocking autophagy via a calpain-dependent mechanism in cells expressing mutant htt or mutant alpha synuclein (Sarkar and Rubinsztein, 2006). Thus increases in endogenous mIns, a precursor for IP3, may indicate neurons and glia are actively trying to prevent autophagy. The increase in mIns observed here and previously in R6/2 (Tkac et al., 2007; Zacharoff et al., 2012) could be a cellular response to limit the autophagy induced by the initial generation of aggregates. Increases in mIns are also reported in early clinical but not preclinical HD (Sturrock et al., 2010; Heikkinen et al., 2012). A similar rise in mIns has been observed early in development of the spinocerebellar ataxias, another CAG repeat disease with intracellular aggregates (Oz et al., 2010, 2011).

Normal htt does not bind to PE. Mutant htt associates with phosphatidylethanolamine, a product of PE and diacylglycerol and a prominent component of CNS membranes (Kegel et al., 2009). Polyglutamine expanded htt also binds PE, albeit to a much lesser extent than IP3 (Kegel et al., 2009). The exact binding site for the mutant htt-phospholipid binding activity is not known. Striatal PE decreased progressively beginning at 6 mo in the Q140s. This occurs early in Q140 (but not BACHD) disease compared to the biochemically measured reductions in phosphoethanolamine and to a lesser extent, ethanolamine reported in stage 3 and 4 HD brains (Ellison et al., 1987). Both studies are in agreement, reporting a loss in basal ganglia but not in cortex. The observed drop in PE concentration may be attributable to mass action effects of mutant htt binding to phosphatidylethanolamine. As such, this metabolite may track the accumulation of mutant htt within cells and the degree of cell membrane disturbances (Kegel et al., 2009).

Cortical taurine concentration was elevated in the cortex, but not the striatum of Q140 mice beginning at 9 mo and not at all in BACHDs. Microdialysis experiments detected elevated extracellular taurine in R6/2 cortex in agreement with the increases observed in total taurine by ^1^H MRS in that model (Behrens et al., 2002; Tkac et al., 2007; Zacharoff et al., 2012). Taurine is synthesized by astrocytes (Reymond *et al*. 1996) and released in response to injury (Huxtable, 1989; Scheller et al., 2000). Tau is an essential brain metabolite involved in multiple CNS functions, including the osmotic regulation, brain development, cytoprotection, neuromodulation, inhibitory functions and the regulation of intracellular calcium homeostasis (Ripps and Shen, 2012). While the functional consequence of the elevated cortical Tau remains unknown, a homeostatic neuroprotective mechanism is possible (Oja and Saransaari, 2011).

The 1C2 antibody recognizes the toxic conformation of the CAG repeat expansion of the mutant huntingtin in both mice and humans (Trottier et al., 1995; Sieradzan et al., 1999; Herndon et al., 2009). The incidence of 1C2 and ubiquitin positive inclusions was comparably high among all R6/2 brains and low among BACHD brains, corroborating the ability of the 1C2 antibody to detect inclusions. These aggregates have also been demonstrated to contain mutant huntingtin (Sieradzan et al., 1999; Herndon et al., 2009). At the end of the R6/2 lifespan, intracellular inclusions, visible by either antibody, were plentiful, consistent with overt disease. By 24 mo of age, intracellular aggregates, visible by either antibody, were hard to find and had not accumulated in BACHD brains. The paucity of 1C2 or ubiquitin positive puncta suggested the cellular pathophysiological process of aggregate formation was just beginning and was not yet widespread. Thus, the absence of cellular pathology in the BACHD brains was consistent with the absence of multiple changes in MRS metabolites.

### Comparison of in vivo ^1^H MRS data in mouse models of HD

Metabolite profiles for the Q140 and BACHD mice detected here mirror the previously reported changes in other mouse models of HD (Table 3). Table 3 arranges the mice left to right along a rough continuum of most to least severe. Prominent among most models are increases in mIns, Gln, and PCr or tCr, while NAA, Glu, GABA and PE decrease. These similarities suggest a metabolic phenotype for fulminant disease. The slower progressing models, represented by the BACHD, present a different pattern of changes indicative of premanifest mouse HD without appreciable degeneration. As catabolic products of cell membrane metabolism, cholines have been interpreted as indicators of neuronal loss(Duarte et al., 2012). Cholines and NAA may indicate more extensive or rapid rates of neurodegeneration. A slower rate of neurodegeneration may provide more time for ordered or phagocytic clearance of cellular debris. Q140 mice may not live long enough for development of sufficient degeneration to trigger alterations in GPC+PC and NAA, as seen in more severe HD mice. In addition, many reported changes only occur at specific ages in longitudinal studies. Such temporal variation in disease progression may reflect homeostatic mechanisms that correct the imbalance detected previously. Data from the Q111 mouse at 6 and 13wk of age, early ages in a mild model, exemplify this (Tkac et al., 2012).

**Table 3.**
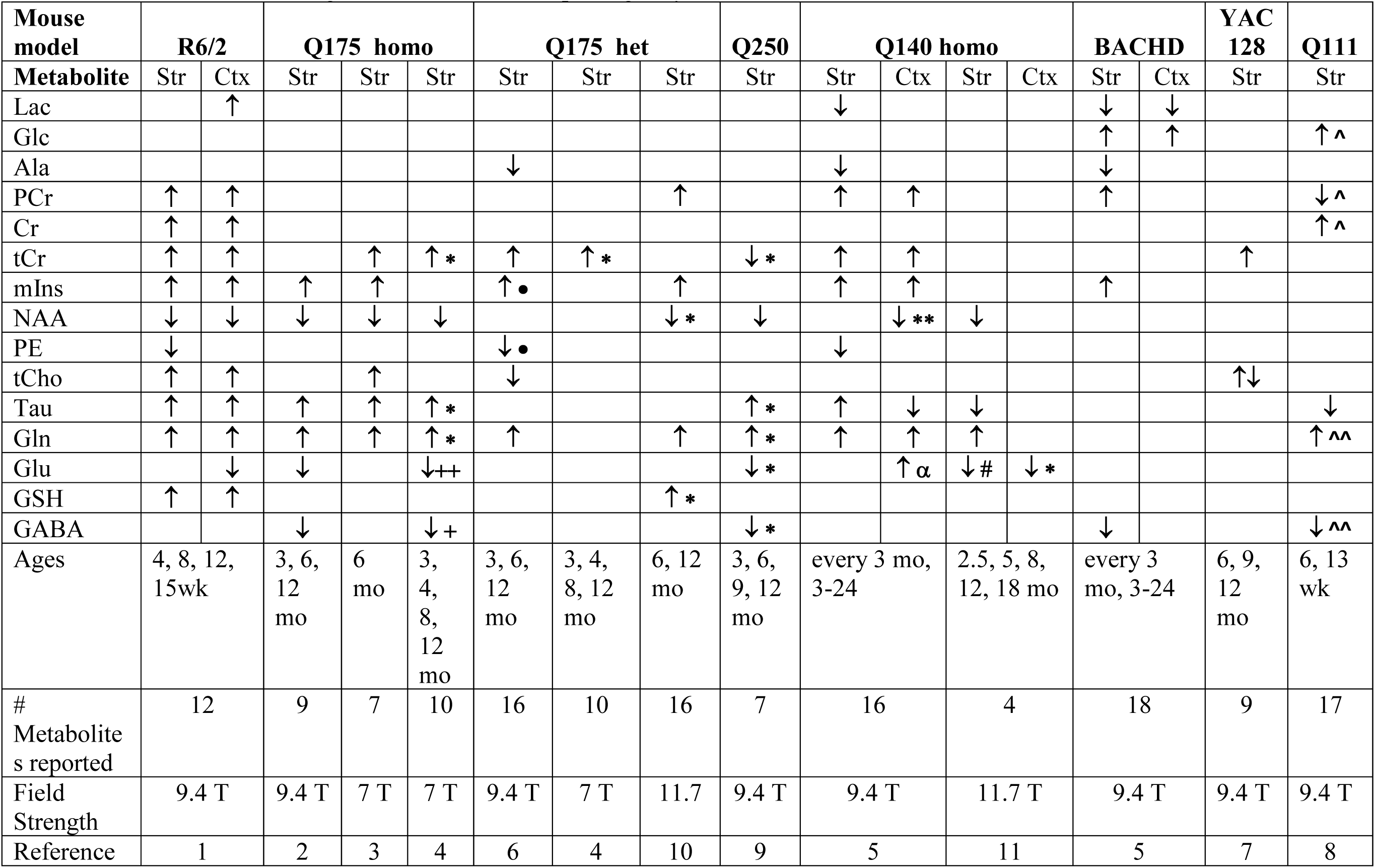
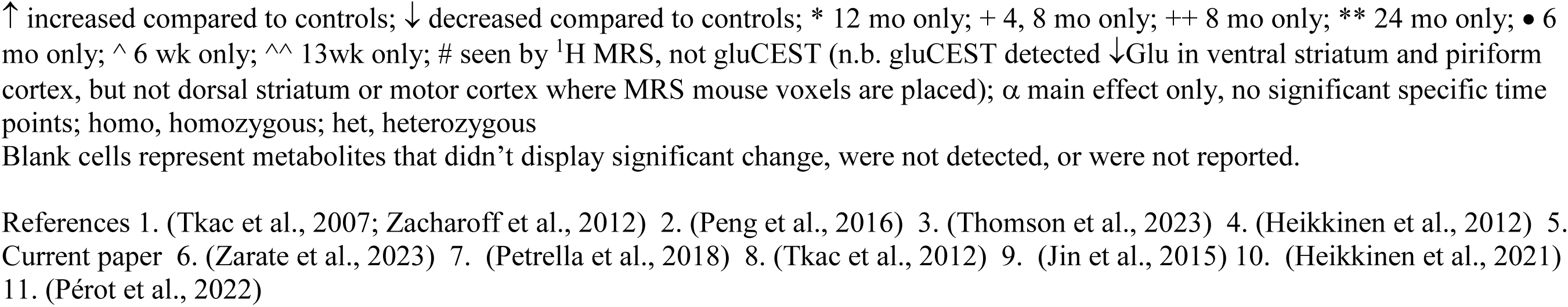
Comparison of HD mouse model significant ^1^H MRS metabolite concentration changes from high field strength studies of dorsal striatum. Low field strength studies or studies reporting only ratioed metabolite values are not included.

Comparison of mouse and human ^1^H MRS data becomes difficult as the mouse studies utilize the most advanced spectroscopy techniques at ultra-high magnetic fields (Mekle et al., 2009; Tkac et al., 2009; Oz and Tkac, 2011), techniques not available for clinical human use (Hoang et al., 1998; Jenkins et al., 1998; Sturrock et al., 2010, 2015; Unschuld et al., 2012; van den Bogaard et al., 2014). In human ^1^H MRS studies, the range of detectable metabolites and the precision of the quantification are limited compared to those in mice (Sturrock et al., 2010, 2015). Even among well executed mouse studies, the difference in resolution obtained from 9.4T and 7T scans may underlie the ability to detect significance in the small changes in a metabolite such as mIns (Heikkinen et al., 2012; Zacharoff et al., 2012). Nevertheless, ^1^H MRS from early stage individuals revealed decreases in NAA and increases in mIns, largely consistent across mouse models scanned to date (Sturrock et al., 2010; Heikkinen et al., 2012; Zacharoff et al., 2012, 2012; Zarate et al., 2023). Mathematical models of metabolite data may yield biomarker measures more sensitive to patterns of change than following concentrations of only a single metabolite (Tsang et al., 2006; Sturrock et al., 2010; Zacharoff et al., 2012).

### Biomarkers

As in prior longitudinal ^1^H MRS analysis of HD mouse models, metabolites were observed to change dynamically with disease progression and lifespan (Tkac et al., 2012). The varying severity and progression timelines among the different HD mice make comparisons among the models difficult. Yet the rough similarity in metabolite changes provide common markers enabling such comparisons. The binary logistic regression models performed here on Q140 ^1^H MRS and other data demonstrate the ability to predict disease in related but genetically distinct HD mice.

Multivariate modeling approaches provide valuable methods for optimizing biomarker construction. Indeed, combinations of functional, anatomical and behavioral biomarkers promise to provide accurate descriptions of disease progression and possibly demonstrate the ability to respond to interventions (Ross et al., 2014; Mason et al., 2018). A machine learning algorithm based upon logistic regression of multiple measures from longitudinal human HD studies can predict risk of progression for participants entering future clinical trials, reducing sample size and expected outcome variability (Koval et al., 2022). Alignment of disease severity by ‘Huntington Age’ (Koval et al., 2022) might reduce some of the variability among both human and mouse HD ^1^H MRS studies (Lowe et al., 2022). Human ^1^H MRS would also benefit from determination by optimal protocols in higher field strength magnets, as used in mice studies. In calculating future combinatorial biomarkers, MRS measures should be included, as changes in metabolite concentrations do not change monotonically and may predate other anatomical or functional measures (Tsang et al., 2006; Tkac et al., 2012; Zacharoff et al., 2012).

## Acknowledgements

The authors would like to thank Drs. Tongbin Li and Gabriel Al-Ghalith for performing PLSDA and exploratory analyses, respectively. This work was generously funded by the CHDI Foundation, Inc.

---

**Supplementary Figure 1.**
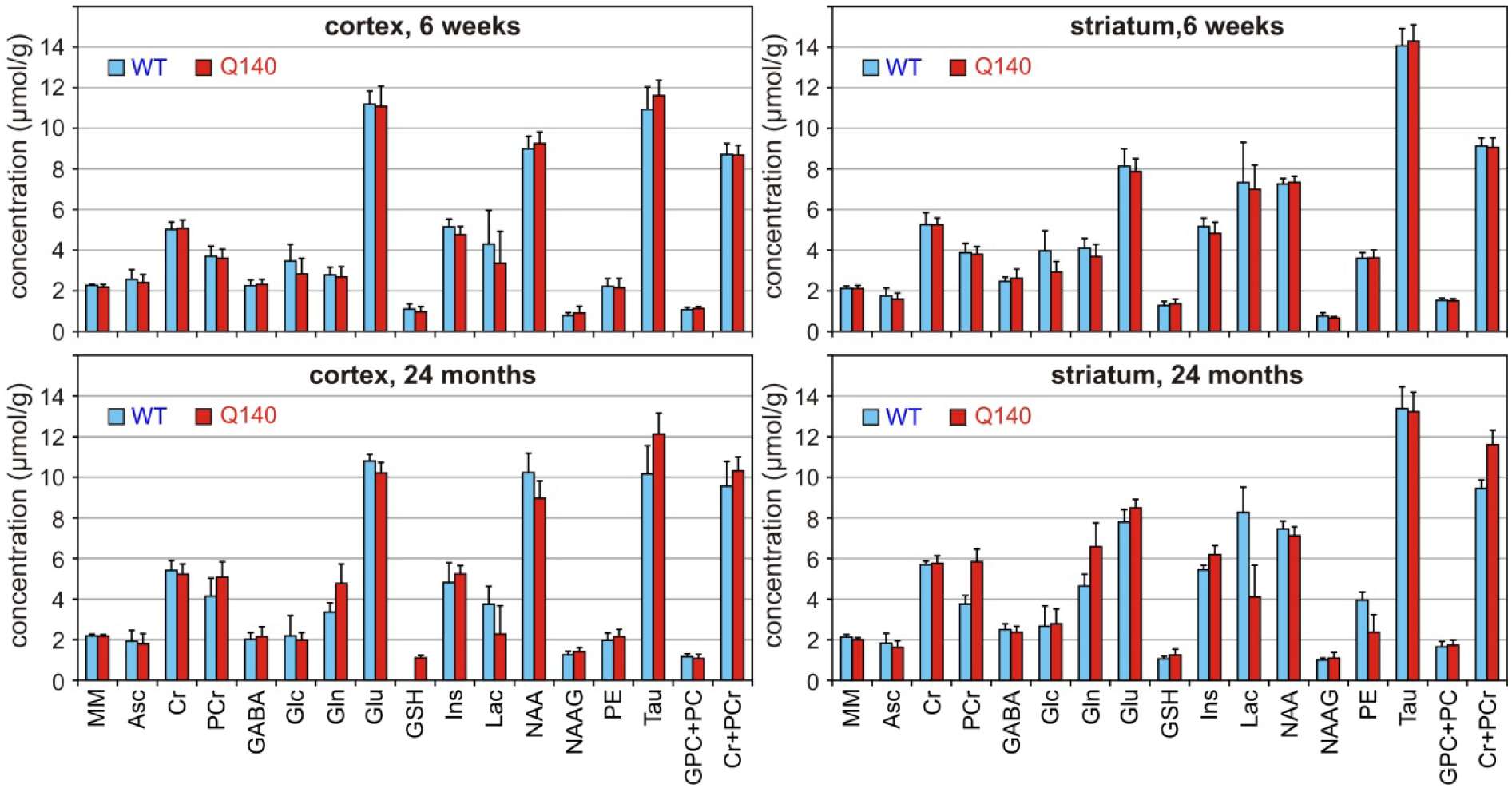
Concentration data from 6 week and 24 month old Q140 and littermate mice striatum and cortex.

**Supplementary Figure 2.**
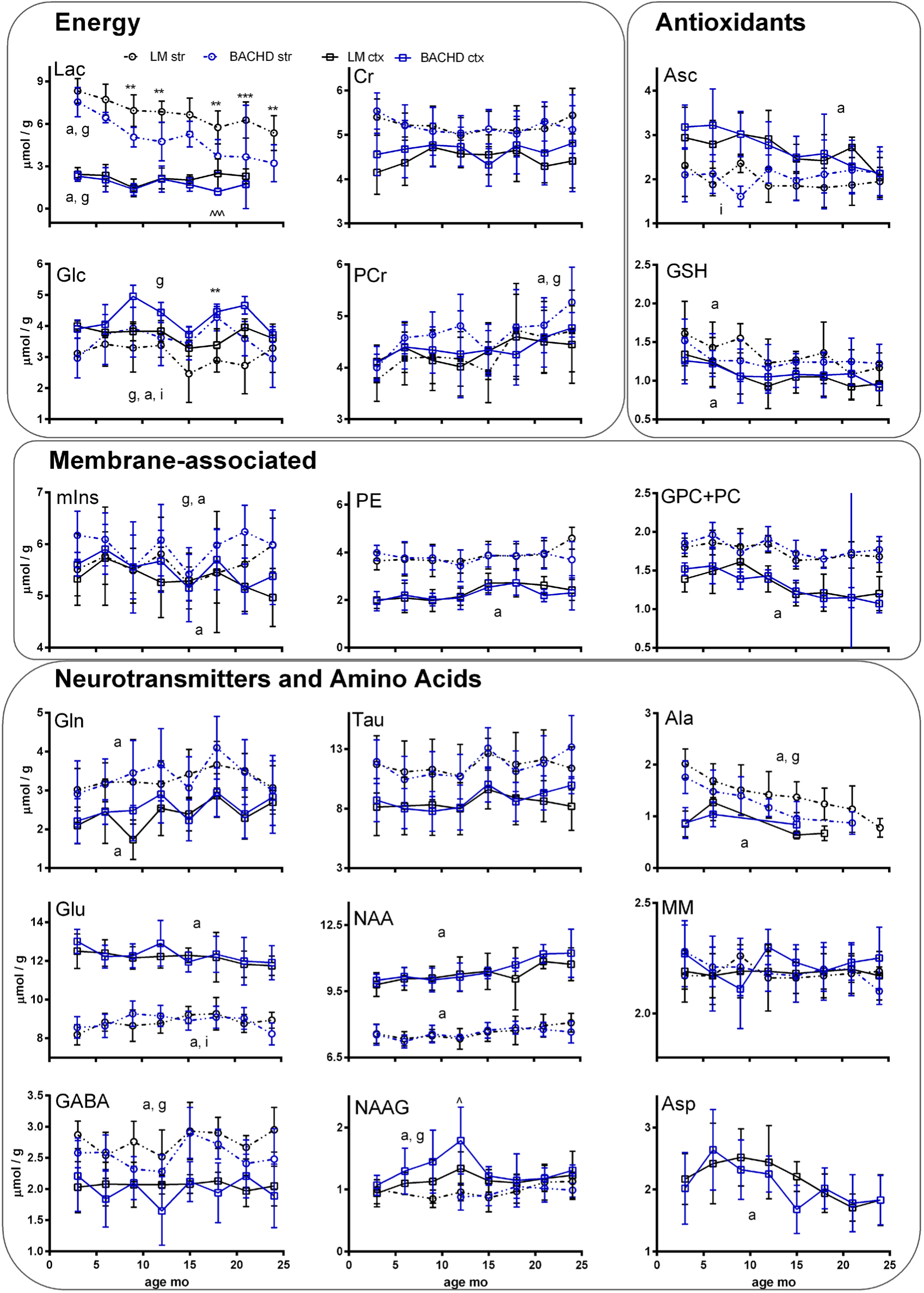
Longitudinal changes in 18 striatal and cortical metabolites in striatum (circles, dotted lines) and cortex (squares, solid lines) over the lifespan of BACHD (blue) and littermate (black) mice. Data are mean ± standard deviation from 8 or more mice of each genotype per time point. Statistical results coded as in Fig. 2.

**Supplementary Fig 3.**
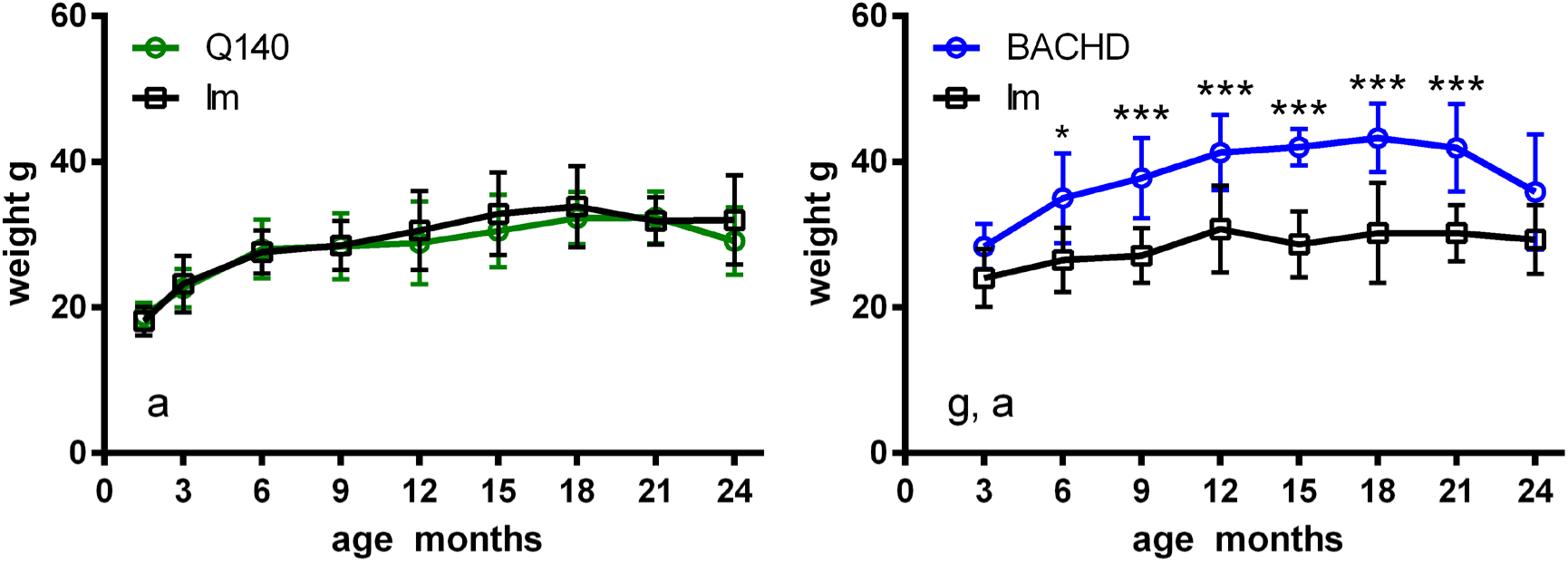
Body weights of Q140 (left, green) and BACHD mice (right, blue). Statistical results coded as in Fig. 2.

**Supplementary Figure 4.**
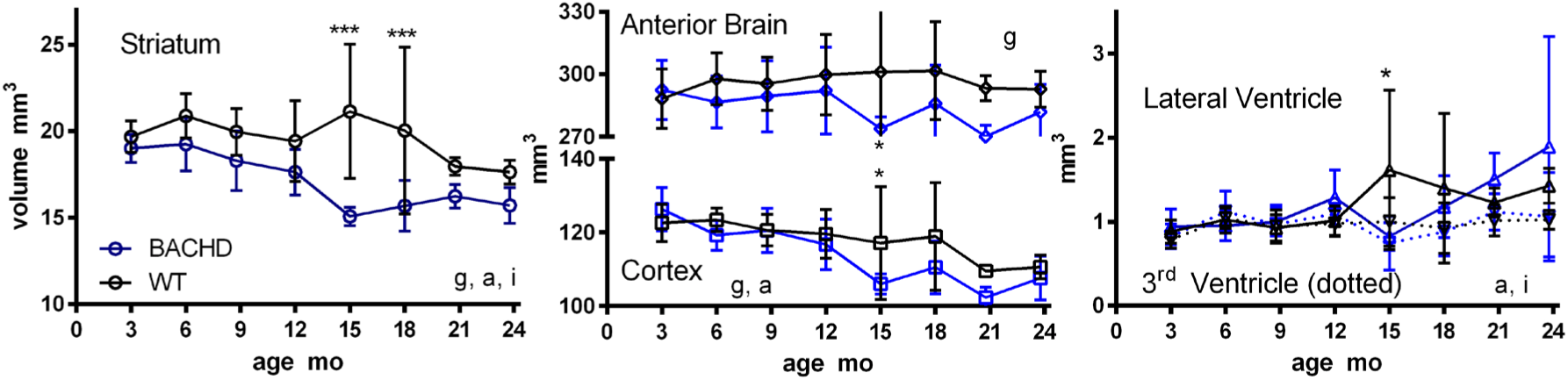
Analysis of regional volume changes across the BACHD (blue) and wild type (black) lifespan as measured in coronal images as illustrated in Fig. 4. Statistical results coded as in Fig. 2. N.B. It should be noted that due to the change in computer consoles used to run the 9.4T magnet, a backup set of BACHD and littermate mice were used for the 15mo time point. The 18 mo and following time points were largely comprised of the original cohort with some backups. Thus the deviance of brain volumes at 15 mo may be attributable to the cohort of mice and not to age or disease.

**Supplementary Figure 5.**
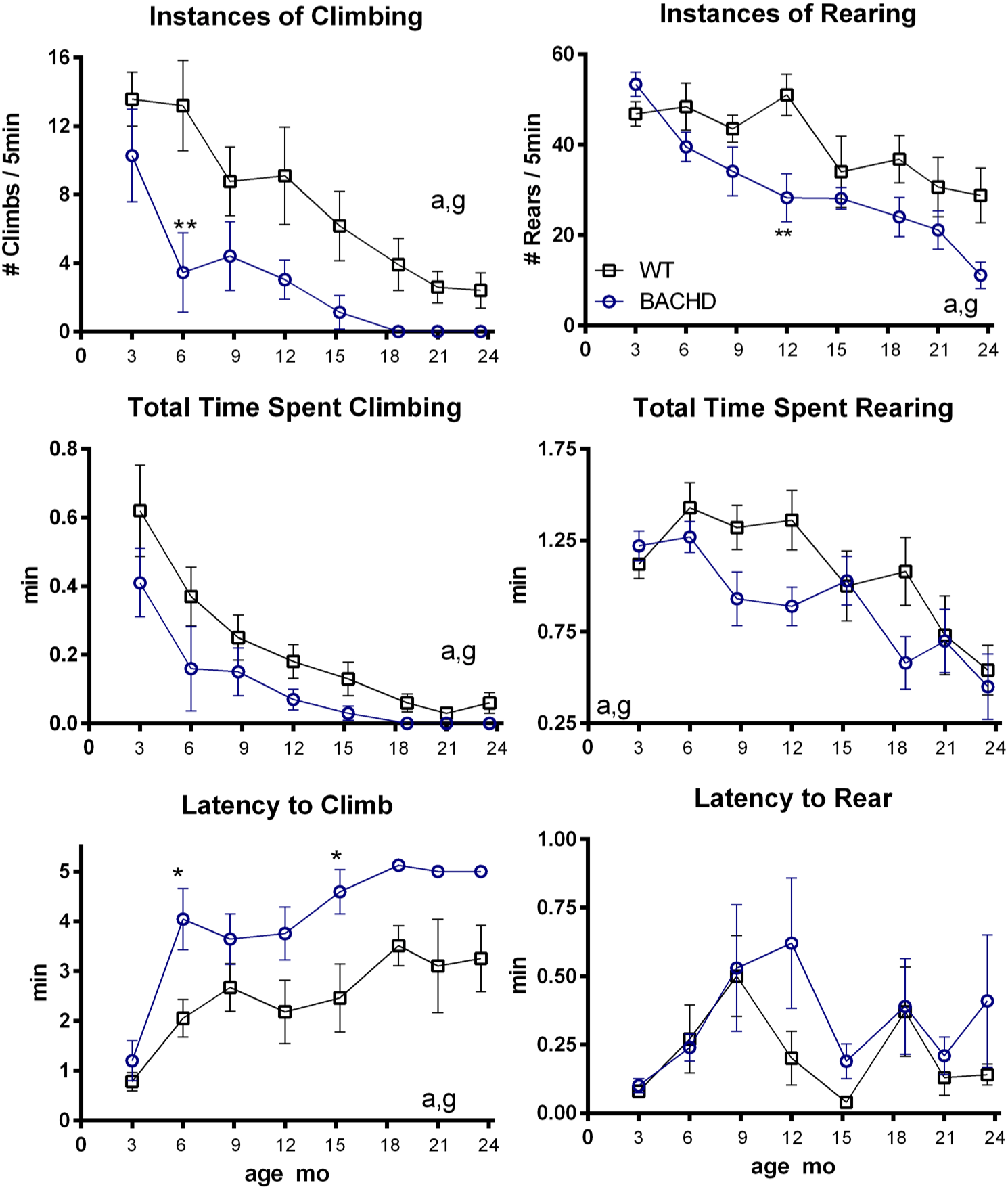
Longitudinal changes in climbing assay behavior over the BACHD (blue circles) and littermate (black squares) lifespan. Statistical results coded as in Fig. 2.

